# Reference-based QUantification Of gene Dispensability (QUOD)

**DOI:** 10.1101/2020.04.28.065714

**Authors:** Katharina Sielemann, Bernd Weisshaar, Boas Pucker

**Author notes:** Email addresses.

## Abstract

**Background:** Dispensability of genes in a phylogenetic lineage, e.g. a species, genus, or higher-level clade, is gaining relevance as most genome sequencing projects move to a pangenome level. Most analyses classify genes as core genes, which are present in all investigated individual genomes, and dispensable genes, which only occur in a single or a few investigated genomes. The binary classification as ‘core’ or ‘dispensable’ is often based on arbitrary cutoffs of presence/absence in the analysed genomes. Even when extended to ‘conditionally dispensable’, this concept still requires the assignment of genes to distinct groups.

**Results:** Here, we present a new method which overcomes this distinct classification by quantifying gene dispensability and present a dedicated tool for reference-based QUantification Of gene Dispensability (QUOD). As a proof of concept, sequence data of 966 *Arabidopsis thaliana* accessions (Ath-966) were processed to calculate a gene-specific dispensability score for each gene based on normalised coverage in read mappings. We validated this score by comparison of highly conserved Benchmarking Universal Single Copy Orthologs (BUSCOs) to all other genes. The average scores of BUSCOs were significantly lower than the scores of non-BUSCOs. Analysis of variation demonstrated lower variation values between replicates of a single accession than between iteratively, randomly selected accessions from the whole dataset Ath-966. Functional investigations revealed defense and antimicrobial response genes among the genes with high-dispensability scores.

**Conclusions:** Instead of classifying a gene as core or dispensable, QUOD assigns a dispensability score to each gene. Hence, QUOD facilitates the identification of candidate dispensable genes, associated with high dispensability scores, which often underlie lineage-specific adaptation to varying environmental conditions.

## Background

Genetic variation is not restricted to single nucleotide polymorphisms or small insertions and deletions but extends also to (large) structural variations. These structural variations include copy number variations (CNVs) and presence/absence variations (PAVs), which can cause substantial variation of the gene content among individual genomes (1,2). The comparative analysis of multiple genomes of the same phylogenetic clade allows the identification of PAVs that are connected to phenotypic traits. In the case of crop species, the identification of PAVs underlying specific agronomic traits which only occur in a single or a few species is feasible (3–5). As more highly contiguous genome sequences become available, pangenomes are suitable to describe and investigate the gene set diversity of a biological clade, e.g. species, genus or higher (6,7).

Genes of a pangenome are thought to be divided into a core and a dispensable gene set, the latter is also often referred to as ‘accessory’ in the literature. Core genes occur in all investigated genomes, whereas dispensable genes only occur in a single or a few genomes (8). In eukaryotic pangenome studies, core and dispensable genes are mostly identified based on sequence similarity e.g. using GET_HOMOLOGUES-EST Markov clustering (9), OrthoMCL gene family clustering (10) or BLASTN (11). Sometimes, a third category of ‘conditionally dispensable’ genes is invoked (12) or genes might be classified as ‘cloud’, ‘shell’, ‘soft-core’ and ‘core’ (13) or even as ‘core’, ‘softcore’, ‘dispensable’ and ‘private’ (14). However, this distinct classification is not based on the biological dispensability of genes and relies on one or multiple arbitrary cutoffs. Some studies consider genes as ‘core’ if these genes occur in at least 90 % of the investigated genomes (11); in other studies, only genes which are found in all genomes are part of the core genome (10). In addition, dependency groups might influence the dispensability of certain genes. The possibility that two genes might be ‘replaced’ by a specific number of other genes has to be considered. Some genes, of e.g. a gene family, might be required in a specific proportion and therefore are only conditionally dispensable (12). Further, assemblies of genomes or transcriptomes might be incomplete leading to artificially missing genes (15). One way to circumvent this is to rely only on high-quality reference genome sequences, thus avoiding additional assemblies which are potential sources of errors.

Here, we present QUOD - a bioinformatic tool to quantify gene dispensability. An *A. thaliana* dataset of about 1,000 accessions was used to calculate a per gene dispensability score derived from the coverage of all genes in the given genomes. This score was validated by comparison of scores of BUSCOs and the functional investigation of genes with high-dispensability scores. Our tool is easy to use for all kinds of plant species. QUOD extends the distinct classification of genes as ‘core’ and ‘dispensable’ based on an arbitrary threshold to a continuous dispensability score.

## Methods

### Selection and preprocessing of datasets

Genomic reads (FASTQ format) of the investigated genomes were retrieved from the Sequence Read Archive (SRA) (16) via fastq-dump. BWA-MEM (v.0.7.13) (17) was applied to map all genomic paired-end Illumina reads to the corresponding reference genome sequence using default parameters as well as *-m* to discard secondary alignments. For *A. thaliana*, all available 1,135 datasets (18) (Additional file 1) were subjected to a mapping against the AthNd-1_v2c genome sequence (19). The resulting BAM files of these mappings were subjected to QUOD.

### Calculation of gene dispensability scores – QUOD

QUOD calculates a reference-based gene dispensability score for each structurally annotated gene based on supplied mapping files (BAM) (one per investigated genome) and a structural annotation of the reference sequence (GFF) (https://github.com/ksielemann/QUOD). The tool is written in Python3 and consists of six different components (Additional file 2). During the first part of the analysis, the read coverage per position (I) as well as the read coverage per gene (II) are calculated. In the next step, genomes with an average coverage below a given cutoff (default=10) are discarded and excluded from further analyses (III). Finally, an input matrix is constructed (IV) and a dispensability score is determined for each gene (V). QUOD assigns high gene dispensability scores to more likely dispensable genes. Optionally, the results can be visualized as a colored histogram and a box plot (VI).

The dispensability score (ds(g)) is calculated as follows (cov.=coverage):

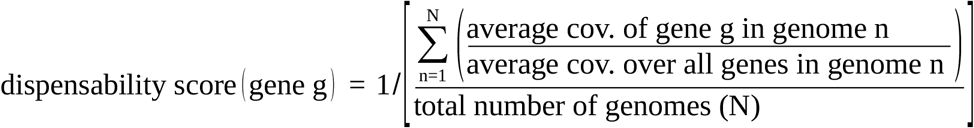

### Comprehension of the dispensability score composition

For further investigation of the score composition of selected genes of interest, the script ‘score_composition.py’ can be used (https://github.com/ksielemann/QUOD/blob/master/score_composition.py). As output, a table including (I) the dispensability score, (II) the average coverage of all investigated genome sequences, (III) the average coverage of the accessions with the highest and (IV) lowest 10 % of all coverage values, respectively, (V) the number of accessions with zero coverage and (VI) the coverage for each accession, separately, is provided. Further, the coverage distribution for each gene can be visualized in a box plot.

### Identification of plastid sequences

Genes of Ath-966 with high similarity to plastid sequences were flagged via BLASTp (20) of the encoded peptides against all organelle peptide sequences obtained from the National Center for Biotechnology Information (NCBI). As a control, the sequences were also searched against themselves. Peptide sequences of Nd-1 with a score ratio ≥ 0.8 were considered plastid-like sequences when comparing BLAST hits against self-hits (19).

### Score comparison between contrasting gene sets

Genes structurally annotated in AthNd-1_v2c were classified with BUSCO v3 (21) running in protein mode on the encoded peptide sequences using ‘brassicales odb10’ (order level) as reference (22). For comparison, BUSCO was additionally executed using ‘chlorophyta odb10’ (phylum level) and ‘embryophyta odb10’ (clade level) as reference. BUSCOs include single-copy genes and universal genes which are present in > 90% of all species in the reference dataset and are used to measure the completeness of assemblies and annotations (21). The scores of BUSCO and non-BUSCO genes were compared using matplotlib (23) for visualization (violin plot) and a Mann–Whitney U test implemented in the Python package dabest (24) for determination of the significance (https://github.com/ksielemann/QUOD/blob/master/BUSCO_comparison.py). Further, a Levene’s test, implemented in the Python package SciPy (25), was calculated to test for equal variances among BUSCO genes and non-BUSCO genes. The dispensability score of non-BUSCO genes might deviate more from the mean as non-BUSCO genes might be less conserved compared to BUSCO genes and might include multi-copy genes. Note that for all analyses performed within this study, the score of the size ‘infinity’ (detected for one gene) was set to the next highest score to enable calculations.

A list of Nd-1 transposable element (TE) genes, which are Nd-1 gene structures overlapping with sequences annotated as TEs, was obtained from Pucker *et al.* (19). First, the score distribution of TE and non-TE genes was determined using a Mann– Whitney U test implemented in the Python package SciPy (25) (https://github.com/ksielemann/QUOD/blob/master/analyse_TE_genes_and_scores.py). Next, the minimal distance of each gene to its closest TE gene was calculated after extracting the gene positions from the Nd-1 annotation file. Mixed linear modelling was performed using Statsmodels v0.12.0 (26) to determine the interaction between the distance to the closest TE gene and the gene dispensability score (https://github.com/ksielemann/QUOD/blob/master/mixed_linear_effects.py).

### Correlation of gene length and exon number with the dispensability score

Length and number of exons per gene were extracted from the Nd-1 annotation file. Linear mixed modelling was performed for gene length, exon number and the gene dispensability score for the whole dataset Ath-966 as well as for three large *A. thaliana* gene families (TAPscan (27)), namely MYBs (28), AP2/EREBP (29) and WRKYs (30) using Statsmodels v0.12.0 (26) (https://github.com/ksielemann/QUOD/blob/master/mixed_linear_effects.py).

### Variation between replicates

A total of 14 genomic datasets of the *A. thaliana* accession Col-0 were received from the SRA (Additional file 3) to assess the technical variation between replicates of the same accession. Col-0 was selected for this analysis, because multiple independent and high-quality datasets are only available for this accession. Each dataset was mapped to the TAIR10 reference genome sequence using BWA-MEM because a Col-0 read mapped against AthNd-1_v2c would result in multiple differences caused by accession-specific differences. The mappings were then subjected to QUOD, expecting a dispensability score close to one for each gene as there should be no variability between datasets of the same accession. As the distributions are different (Kolmogorov-Smirnov test, p ≈ 3e-27) and the sample size (n) is high, the Levene’s test was selected to test for equal variances, regarding the gene dispensability scores. The test was applied for (1) the dataset including replicates only and (2) iteratively (100x), randomly chosen subsets (n=14) of Ath-966 (https://github.com/ksielemann/QUOD/blob/master/variance_in_repl_test.py).

### Functional annotation

All genes of the *A. thaliana* Nd-1 genome sequence were annotated via reciprocal best blast hits (RBHs) and best BLAST hits against Araport11 (19). Functional enrichment analyses (PANTHER protein classes and ‘biological process’ GO terms) were performed using the PANTHER Classification System of the Gene Ontology (31).

### Read mapper comparison

To evaluate the impact of the read mapping, the results of different mappers were compared. In addition to BWA-MEM (v.0.7.13; see above) (17), Bowtie2 (v2.4.1; default parameters) (32) and STAR (v2.5.1b) (33) were selected for this analysis. STAR parameters required alignments with a similarity of at least 95% over at least 90% of the read pair length. The average coverage values per gene were investigated for correlation using the Spearman correlation coefficient implemented in the Python package SciPy (25).

### Data Availability

The tool QUOD (QUOD.py) can be downloaded from GitHub (https://github.com/ksielemann/QUOD; http://doi.org/10.5281/zenodo.4066818). A data set to test QUOD is available on ‘PUB - Publications at Bielefeld University’ (http://doi.org/10.4119/unibi/2946079).

## Results

In this study, a bioinformatic tool was developed to calculate a gene-specific dispensability score based on the normalised coverage in a read mapping. QUOD allows the quantification of dispensability by calculation of a single score for each gene (Figure 1). The binary classification of gene dispensability can be compared to the original method of mRNA detection by endpoint RT-PCR providing only qualitative results (34–36) which was replaced by quantitative analyses like RNA-Seq.

**Figure 1:**
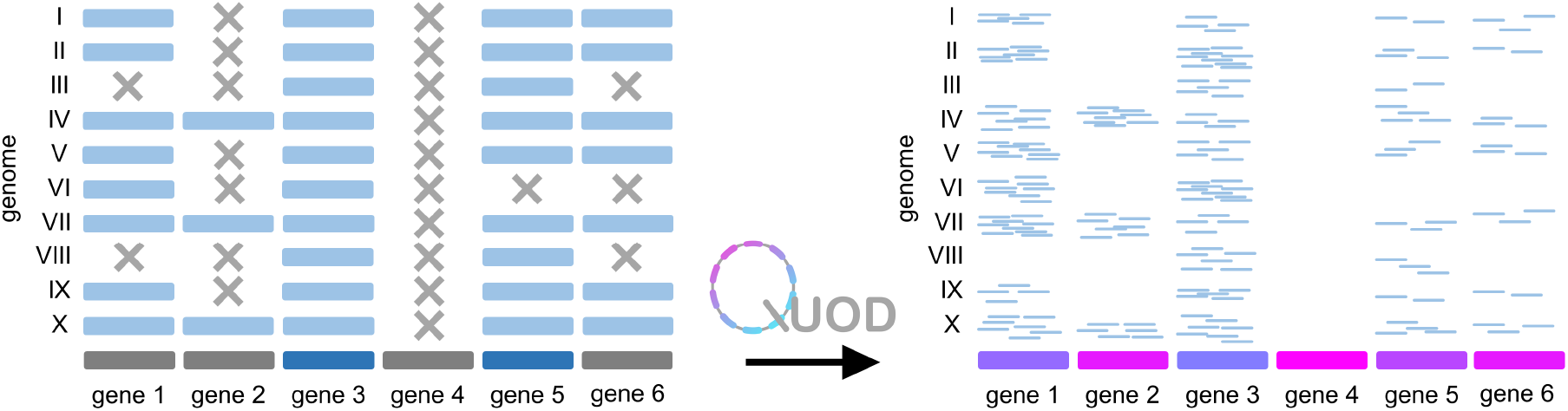
Illustration of the QUOD method using a fictional dataset. On the left side, genes are classified as ‘core’ (dark blue) or ‘dispensable’ (dark grey) according to a cutoff. On the right side, gene dispensability is quantified according to a dispensability score based on the normalised coverage in a read mapping (I-X: investigated genomes). Coloring of genes (right side) indicates different dispensability scores. Extremely rare genes, which are absent from most genomes but present in the reference, can be easily detected using QUOD.

### Gene dispensability scores

The gene dispensability score would initially be dependent on the sequencing depth per genome. By division of the average coverage of gene g in genome n (N = total number of investigated genome sequences) by the average coverage over all genes in genome n, the score is normalised for differences in the sequencing depth of the investigated genomes. A high value indicates that a gene is likely to be missing in some genomes and therefore more likely dispensable than a gene with a lower dispensability score. Due to this quantification approach, this method is not based on an arbitrary cutoff to determine the core genome and the dispensable genome of any given pangenome dataset. An example: Using a cutoff of ‘gene n occurs in at least 90 % of all genomes’ to be considered a ‘core’ gene (dark blue), genes 1,2,4 and 6 (dark grey) would be considered ‘dispensable’ (Figure 1). However, considering the coverage (right panel), it is not clear if e.g. gene 1 is truly biologically dispensable. QUOD does not rely on any thresholds for the classification of genes into ‘core’ and ‘dispensable’, but provides a score based on the normalised coverage in a read mapping. The genes could theoretically be ranked as well using the percentage of presence/absence of a gene in the investigated genomes. However, this alternative approach would still rely on a threshold, e.g. the number of mapped reads for a gene to be considered present in a genome. This threshold is avoided using the QUOD method.

As a proof of concept, *A. thaliana* sequence reads of 1,135 accessions were mapped to the *A. thaliana* Nd-1 genome sequence. All accessions with less than 10-fold read coverage were discarded. The remaining sequencing dataset Ath-966 was analysed with QUOD to calculate a dispensability score for each gene (Figure 2). Genes with high dispensability scores, colored in pink, are considered to be likely dispensable, whereas genes with dispensability scores close to one (dark purple/dark blue) are considered to be core genes.

**Figure 2:**
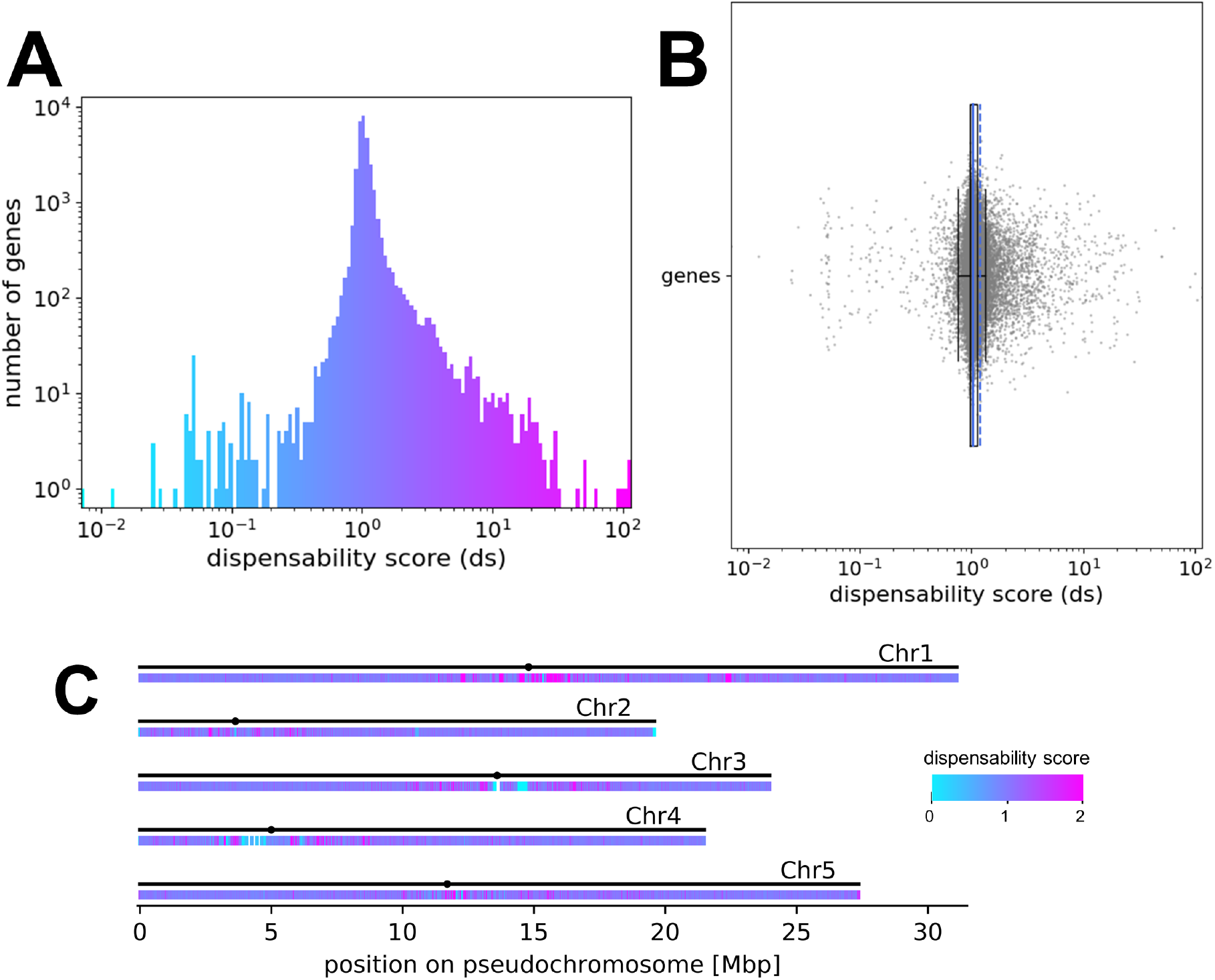
Distribution of the gene dispensability scores for Ath-966. A) Histogram coloured according to the dispensability score. The x-axis represents the dispensability score and the y-axis shows the number of genes in each bin in logarithmic scale. B) Box plot representing the dispensability score (x-axis) of all genes (y-axis). The mean is represented by the dashed blue line, the other blue line represents the median of the scores. C) Genome-wide distribution of genes with different dispensability scores in *A. thaliana* Nd-1. The coloured heatmap shows the respective gene dispensability scores. There are low (blue) and high (pink) scoring genes clustered in repetitive regions, including centromeric and telomeric areas. The x-axis represents the size (in Mbp) of each pseudochromosome in the assembly. The black dots represent the position of the centromeres of the five chromosomes in the AthNd1_v2c assembly (19).

### Genome-wide distribution of the gene dispensability scores

Next, the genome-wide distribution of genes with specific gene dispensability scores was investigated in *A. thaliana* (Figure 2C). A high plasticity between accessions, which means a high number of genes with exceptionally high and low scores (pink and blue), in the (peri-)centromeric regions is visible based on a heatmap (Figure 2C).

As high and low scoring genes cluster in repetitive regions (mainly centromeres), the score distribution of TE genes was investigated (Additional file 4). Scores of TE genes are evenly distributed across all dispensability scores. In total, the mean score of TE genes (mean ds ≈ 1.501) is significantly higher when compared to non-TE genes (mean ds ≈ 1.168) (Mann-Whitney U test, p ≈ 6E-8), which are more frequent across scores close to one. Moreover, the minimal distance of each gene to its closest TE gene and the dispensability scores revealed no relation (Additional file 4).

To test the hypothesis whether genes with higher dispensability scores/more likely dispensable genes are shorter and whether introns accumulate in core genes, the correlation of the gene dispensability score with gene length and exon number, respectively, were determined for the Ath-966 and for three selected gene families separately. However, no clear trend was detectable (Additional file 5).

### Validation of the reliability

Validation of the reliability of the gene dispensability quantification was achieved by comparison of BUSCOs and non-BUSCOs (Additional file 6). BUSCO genes show on average slightly lower scores than non-BUSCO genes for all three reference datasets (p < 0.001, Mann-Whitney U test). Levene’s test was used to test for equal variances. The results show that the variances for all reference datasets differ significantly between BUSCO and non-BUSCO genes (p < 0.001, Levene’s test). Thus, the deviation of the dispensability score from the respective mean is significantly higher for non-BUSCO genes in comparison to BUSCO genes.

Further, functional annotation of BUSCO outliers, which are genes of the ‘brassicales odb10’ BUSCO gene set with dispensability scores below 0.75 or above 1.25, revealed, amongst others, several repeat proteins, transmembrane proteins, a ‘stress induced protein’, and multiple hypothetical proteins (Additional file 7).

Genes with high and low gene dispensability scores were assessed in more detail. Among genes with high dispensability scores, several significantly enriched PANTHER protein classes were detected, e.g. defense/immunity and antimicrobial response proteins, small GTPases and G-proteins (Table 1). Among genes with dispensability scores < 0.8, genes encoding proteins of the extracellular matrix were significantly enriched (Table 1). ‘Biological process’ GO term enrichment revealed several significantly enriched terms associated with the regulation of cellular processes as well as associated with response to stimuli among genes with dispensability scores > 2 (Table 1). Genes with low dispensability scores show enrichment of primary metabolic processes (Table 1).

**Table 1:** Closer investigation of genes with scores >2 and genes with scores < 0.8. Significantly enriched PANTHER protein classes (padj < 0.05) as well as significantly enriched GO biological process terms (padj < 0.05) are shown. Abbreviations: p = process, mp = metabolic process.

| **PANTHER protein classes (padj < 0.05) of genes with scores >2** |  |
| --- | --- |
| small GTPase (PC00208) | 4.21E-05 |
| defense/immunity protein (PC00090) | 4.24E-05 |
| antimicrobial response protein (PC00051) | 5.24E-05 |
| G-protein (PC00020) | 4.05E-04 |
| protein class (PC00000) | 2.04E-03 |
| Unclassified | 2.44E-03 |
| protein-binding activity modulator (PC00095) | 3.72E-02 |
| **PANTHER protein classes (padj < 0.05) of genes with scores <0.8** |  |
| extracellular matrix structural protein (PC00103) | 5.40E-06 |
| extracellular matrix protein (PC00102) | 1.14E-05 |
| Unclassified | 2.68E-05 |
| protein class (PC00000) | 3.57E-05 |
| metabolite interconversion enzyme (PC00262) | 3.04E-02 |
| **GO biological process terms (padj < 0.05) of genes with scores >2** |  |
| cellular p (GO:0009987) | 2.62E-08 |
| mp (GO:0008152) | 4.62E-07 |
| cellular mp (GO:0044237) | 2.85E-06 |
| primary mp (GO:0044238) | 2.37E-05 |
| organic substance mp (GO:0071704) | 3.02E-05 |
| regulation of cellular mp (GO:0031323) | 9.82E-04 |
| regulation of biosynthetic p (GO:0009889) | 9.92E-04 |
| regulation of cellular biosynthetic p (GO:0031326) | 1.04E-03 |
| regulation of cellular macromolecule biosynthetic p (GO:2000112) | 2.27E-03 |
| regulation of macromolecule biosynthetic p (GO:0010556) | 2.52E-03 |
| regulation of primary mp (GO:0080090) | 2.90E-03 |
| macromolecule mp (GO:0043170) | 2.93E-03 |
| regulation of nitrogen compound mp (GO:0051171) | 4.53E-03 |
| regulation of RNA mp (GO:0051252) | 4.69E-03 |
| positive regulation of biological p (GO:0048518) | 4.89E-03 |
| response to organic substance (GO:0010033) | 4.91E-03 |
| positive regulation of cellular p (GO:0048522) | 6.62E-03 |
| regulation of RNA biosynthetic p (GO:2001141) | 6.67E-03 |
| regulation of mp (GO:0019222) | 6.72E-03 |
| regulation of nucleobase-containing compound mp (GO:0019219) | 6.74E-03 |
| regulation of nucleic acid-templated transcription (GO:1903506) | 6.95E-03 |
| developmental p (GO:0032502) | 7.01E-03 |
| response to hormone (GO:0009725) | 7.25E-03 |
| regulation of transcription, DNA-templated (GO:0006355) | 7.27E-03 |
| response to oxygen-containing compound (GO:1901700) | 7.53E-03 |
| anatomical structure development (GO:0048856) | 7.62E-03 |
| nitrogen compound mp (GO:0006807) | 1.25E-02 |
| response to endogenous stimulus (GO:0009719) | 1.48E-02 |
| regulation of gene expression (GO:0010468) | 2.91E-02 |
| system development (GO:0048731) | 3.44E-02 |
| regulation of macromolecule mp (GO:0060255) | 3.45E-02 |
| cellular lipid mp (GO:0044255) | 4.10E-02 |
| clathrin coat disassembly (GO:0072318) | 4.14E-02 |
| multicellular organismal p (GO:0032501) | 4.19E-02 |
| vesicle uncoating (GO:0072319) | 4.26E-02 |
| **GO biological process terms (padj < 0.05) of genes with scores <0.8** |  |
| cellular p (GO:0009987) | 6.35E-07 |
| mp (GO:0008152) | 1.35E-06 |
| organic substance mp (GO:0071704) | 8.49E-06 |
| cellular mp (GO:0044237) | 2.92E-05 |
| nitrogen compound mp (GO:0006807) | 5.35E-04 |
| primary mp (GO:0044238) | 5.76E-04 |
| macromolecule mp (GO:0043170) | 3.82E-03 |
| organonitrogen compound mp (GO:1901564) | 9.67E-03 |
| localization (GO:0051179) | 4.87E-02 |

The function of the 100 genes with the highest gene dispensability scores was examined in detail for Ath-966 (Additional file 8). Fourteen genes of Ath-966 are annotated as “disease resistance proteins”, whereas seven genes are annotated as transposons/transposases. Four genes are described as hypothetical proteins and 24 genes have no functional annotation. In addition, an example for lineage specific adaptation is provided (Additional file 9). The gene NdCChr1.g3308 has a dispensability score of approx. 10. For 870 accessions, which account for approx. 90 % of Ath-966, no coverage was detected. The gene is annotated as resistance gene mediating resistance against the bacterial pathogen *Pseudomonas syringae*.

Next, the variation between replicates of the same accession (Col-0) was determined (Additional file 10). The variation of the gene dispensability score distribution of the replicate dataset (one accession) (σ^2^ ≈ 0.0226) is significantly lower than the variation between all iteratively, randomly selected subsets of *A. thaliana* accessions (σ^2^ ≈ 0.0392) (Levene’s test, p ≈ 4e-19). The average coverage per gene using different read mappers revealed strong correlations in all comparisons (Additional file 11). The coverage correlations, calculated using Spearman correlation coefficient, between BWA-MEM and bowtie2 (r ≈ 0.810, p ≈ 0.0), BWA-MEM and STAR (r ≈ 0.814, p ≈ 0.0) as well as bowtie2 and STAR (r ≈ 0.760, p ≈ 0.0) are similar.

## Discussion

QUOD was developed for the quantification of gene dispensability in plant pangenome datasets. Multiple accessions of several plant species have been sequenced and pose potential use cases for QUOD (Additional file 12). Dropping sequencing costs will lead to an increasing availability of comprehensive sequence datasets which would permit the application of QUOD. Additionally, QUOD is not restricted to plants, but could be applied to other species (e.g. pig (37)). However, an accurate determination of gene dispensability scores free of systematic biases might rely on a uniform selection of genomes from the respective taxonomic group and on uniform read coverage of genes. In addition, non-random fragmentation of DNA prior to sequencing (38) may cause biases. The variation among replicates of the same accession (Col-0; σ^2^ ≈ 0.0226) might be attributed to technical biases, e.g. during sequencing library preparation. The comparison of different read mappers revealed a significant correlation for the average coverage per gene. Outlier samples, detected by the investigation of the average coverage per gene using different read mappers, might indicate technical issues. Even though the correlations are strong, the same tool with the same parameter settings needs to be used for the read mapping of all compared datasets within one single QUOD run.

Most genes show dispensability scores close to one as the majority of genes are widespread across species. The aim of QUOD is mainly the identification of the ‘outliers’ and therefore the more dispensable genes, which are genes not present in all genomes. These dispensable genes represent a smaller fraction of the genome than the core genes. Genome level patterns are expected to be similar for all species. Further, QUOD is not an alternative to PAV detection methods as groups of genes can still always be defined using PAV methods, but QUOD provides a quantitative measurement for these cases.

As already stated in the Introduction, genome assemblies might be incomplete leading to artificially missing genes (15). One way to circumvent this is to rely on a high-quality reference genome sequence, thus avoiding additional assemblies which are potential sources of errors. Recently released telomere-to-telomere assemblies indicate that these resources will be available for many plant species in the near future (39). Further, the usage of QUOD with a synthetic reference derived from multiple assemblies is possible and can be implemented in the future. A graph-based assembly of a pangenome comprising multiple accessions is already feasible for bacteria (40–42). However, for large plant genome sequences graph-based pangenome assembly is computationally expensive and not yet robust for complex structural variants like inversions(43). Even though there are still several shortcomings, like loss of the sample information (44), improved methods might be available in the near future and could be used for the improved quantification of gene dispensability.

### Genome-wide distribution of the gene dispensability scores

The genome-wide distribution of all gene dispensability scores (not only BUSCO genes) of the *A. thaliana* genomes reveals the origin of exceptionally low dispensability scores (Figure 2). Low scoring genes, which are colored in light blue in Figure 2, might be TEs and other repeat genes associated with collapsed sequences in the assembly. An accurate determination of the dispensability scores of these genes might be possible using ideal genome sequences without any collapsed regions and with specific read mappings e.g. using high quality long reads. However, low scoring genes could still be useful to determine amplified TEs and other repeat genes. Moreover, the genome-wide distribution plot (Figure 2C) shows that high and low scoring genes cluster in repetitive regions, like centromeres or telomeres. Very similar sequences, e.g. members of a gene family or close paralogs, might cause read mapping errors confounding biases in the dispensability scores of these genes.

Additionally, this can be explained by variation in the recombination rate (45) and active TEs in these regions. It was previously proposed, that dispensable genes are likely located closer to TEs which are important factors in genome evolution (9). However, in the results of our study, TE genes are widely distributed across all dispensability scores as TEs can occur with variable copy numbers in genomes leading to low scores and can as well be dispensable. Other studies detected a high number of TEs in the dispensable genome (46). However, it is possible that only certain TE families might be truly dispensable. One limitation is the accurate assignment of reads to repetitive sections of the reference sequence during the read mapping (15). Further, only a fraction of transposons might be correctly assembled and annotated due to several computational challenges in highly repetitive and peri-centromeric regions (47). Therefore, a different strategy might be needed to accurately quantify dispensability of TEs. A high quality annotation of transposons and a following exclusion of these genes from the analysis or improved read mapping to the consensus sequence might improve the results. Again, long reads could be an alternative solution to handle regions which might be ambiguous in read mappings. Moreover, heterochromatin or genome-purging mechanisms (48) could influence the gene dispensability scores in these regions.

Additionally some of the low scoring genes were flagged as plastid-like sequences as original sequencing data from plants contain high amounts of reads originating from plastid sequences (49,50). Biases due to this plastid read contamination inflate the coverage of sequences with high similarity to plastid sequences, resulting in an exceptionally low gene dispensability score.

### Validation of the reliability

We validated the reliability of the gene dispensability score by showing that more conserved BUSCO genes get significantly lower dispensability scores than non-BUSCO genes (Additional file 6). Based on the distribution of the scores in the violin plot (Additional file 6), the difference between BUSCOs and non-BUSCOs appears small, even though the difference is significant (U test, p ≈ 4E-113, brassicales reference). It is important to note that non-BUSCO genes can be highly conserved. Consequently, the difference is only visible at the group level. The difference in the dispensability scores of BUSCOs and non-BUSCOs is low as expected, because conserved multiple-copy genes are not included in the BUSCO gene set (21). Therefore, the variance of the dispensability scores of non-BUSCO genes is significantly larger than the variance among BUSCO genes: non-BUSCO genes comprise highly conserved multi-copy genes as well as less conserved genes. Further, functional annotation of BUSCO outliers revealed several repeat proteins and transmembrane proteins. Repeat proteins might lead to read mapping errors and consequently artificial variations in coverage and dispensability scores. Transmembrane proteins are thought to be involved in biotic stress response and might not be essential for some accessions and therefore dispensable (51). This could explain the absence in some genomes resulting in high dispensability scores of these genes. Therefore, many important, lower-scoring genes might lie outside of the BUSCO reference set.

Functional annotation of the 100 most likely dispensable genes revealed a high number of uncharacterised proteins, disease resistance proteins as well as transposons and transposases in the *A. thaliana* genomes. It is possible that these genes are undergoing pseudogenization and have not been functionally annotated due to the lack of a visible phenotype when mutated. TEs were detected in other studies as contributors to large structural variations between species and individuals and considered as a substantial part of the dispensable genome (46). Previous pangenome analyses also revealed that the dispensable genome comprises functions like ‘defense response’, ‘diseases resistance’, ‘flowering time’ and ‘adaptation to biotic and abiotic stress’ (9,11,13). Comparable results were detected for the enriched protein classes and ‘biological process’ GO terms (Table 1), even though very general terms, like ‘protein class’, give little evidence about the function of genes. Moreover, we provide a specific example for lineage specific adaptation associated with a high dispensability score (Additional file 9): a gene mediating resistance against the bacterial pathogen *Pseudomonas syringae*. Therefore, in depth investigation of genes with high dispensability scores can result in the identification and characterization of phenotypic variation (52) and important agronomic traits (13). We envision several applications for the gene dispensability score generated by QUOD: (1) more accurate prediction if a gene is associated with a specific trait, (2) development of dependency gene networks, and (3) improved modeling of the evolutionary value of genes.

## Conclusions

QUOD (reference-based QUantification Of gene Dispensability) overcomes the problem of labeling genes as ‘core’ or ‘dispensable’ through implementation of a quantification approach. Instead of a distinct classification, QUOD provides a ranking of all genes based on assigned gene-specific dispensability scores and therefore does not rely on any thresholds.

## Declarations

### Ethics approval and consent to participate

Not applicable.

### Consent for publication

Not applicable.

### Availability of data and materials

The tool QUOD for the reference-based QUantification Of gene Dispensability (QUOD.py) can be downloaded from GitHub (https://github.com/ksielemann/QUOD; http://doi.org/10.5281/zenodo.4066818).

### Authors’ contributions

KS, BW and BP designed the study, performed the experiments, analysed the data, and wrote the manuscript. All authors read and approved the final version of this manuscript.

## Acknowledgements

We thank members of Genetics and Genomics of Plants for discussion of preliminary results. We are very grateful to Janik Sielemann and Nathanael Walker-Hale for helpful comments on the manuscript. We acknowledge support for the Article Processing Charge by the Deutsche Forschungsgemeinschaft and the Open Access Publication Fund of Bielefeld University. We thank the CeBiTec Bioinformatic Resource Facility team for great technical support.

## Additional files

Additional file 1 (.tsv): SRA IDs of datasets downloaded to conduct the QUOD analysis of the *A. thaliana* genomes.

Additional file 2 (.pdf): Illustration of the different components of QUOD.

Additional file 3 (.tsv): SRA/ENA IDs of datasets downloaded to conduct the analysis of replicates (Col-0).

Additional file 4 (.pdf): Distribution of scores of TE genes and non-TE genes and correlation of the distance to the closest TE gene with the gene dispensability score of the *A. thaliana* genomes.

Additional file 5 (.pdf): Correlation of gene length and exon number with the dispensability scores of the *A. thaliana* genomes.

Additional file 6 (.pdf): Comparison of BUSCO analyses for ‘chlorophyta’, ‘brassicales’ and ‘embryophyta’ as reference.

Additional file 7 (.tsv): Functional annotation of BUSCO outliers (using ‘brassicales odb10’ as reference) with a dispensability score smaller than 0.75 or greater than 1.25.

Additional file 8 (.tsv): Functional annotation of the 100 most likely dispensable genes of the *A. thaliana* genomes.

Additional file 9 (.pdf): Example for lineage specific adaptation.

Additional file 10 (.pdf): Analysis of variance of the gene dispensability score calculated for replicates of the *A. thaliana* Col-0 accession and iteratively, randomly chosen subsets of the whole dataset Ath-966.

Additional file 11 (.pdf): Correlation of the average coverage per gene using three different read mappers: BWA-MEM, bowtie2 and STAR.

Additional file 12 (.tsv): Examples of diploid species where multiple cultivars were already sequenced.

