## Additional file 2 for "Reference-based QUantification Of gene Dispensability (QUOD)"

**Input**

mapping files (BAM)
(one per investigated genome)

structural annotation of the
reference sequence (GFF)

↓

Calculate read coverage per position (I)

↓

Calculate read coverage per gene (II)

↓

Discard genomes with an average coverage below a given cutoff (default=10) (III)

↓

Construct input matrix (IV)

↓

Determine dispensability score for each gene (V)

$$\text{dispensability score (gene g)} = 1 \Big/ \left[ \frac{\sum_{n=1}^{N} \left( \frac{\text{average cov. of gene g in genome n}}{\text{average cov. over all genes in genome n}} \right)}{\text{total number of genomes (N)}} \right]$$

↓

Visualize the score distribution (VI)
