## Additional file 4 for "Reference-based QUantification Of gene Dispensability (QUOD)"

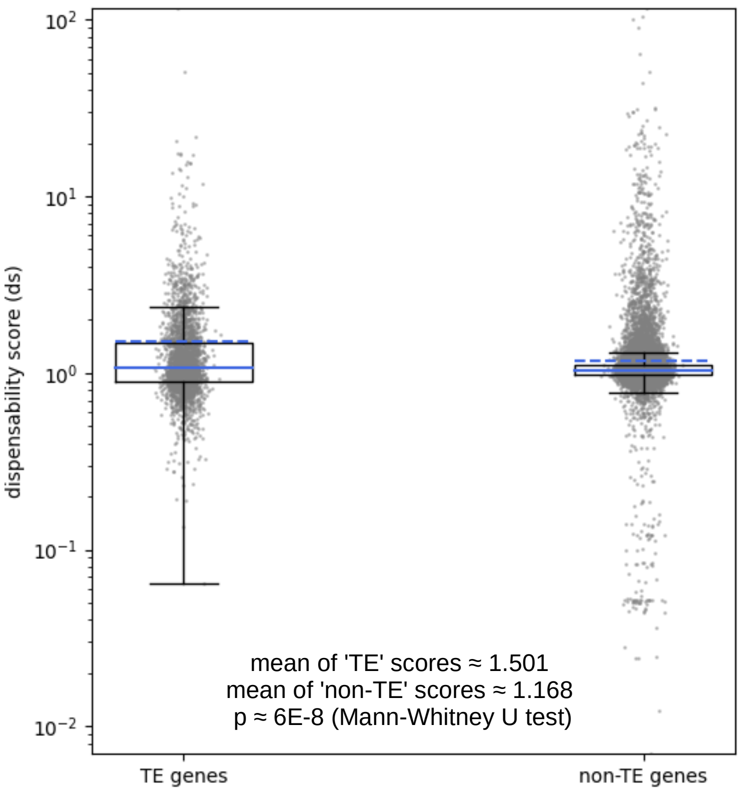


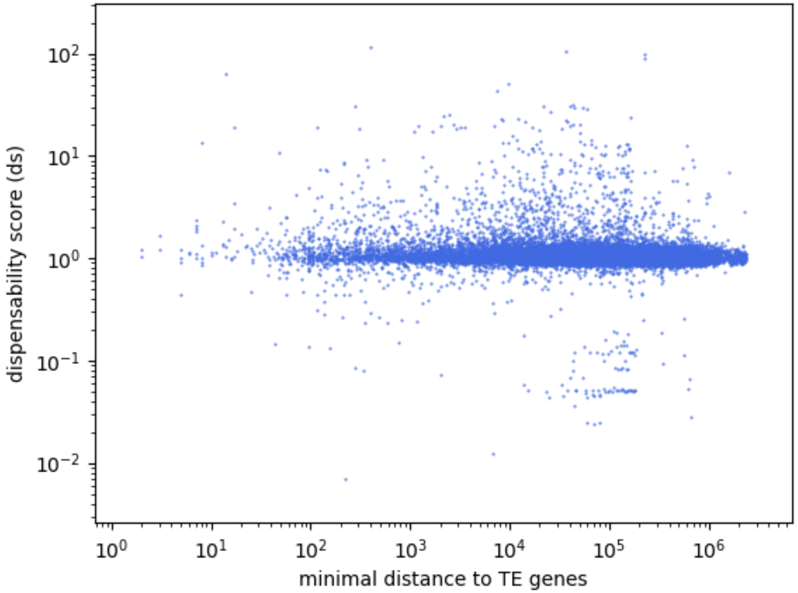


```
                 Mixed Linear Model Regression Results
=====================================================
Model:            MixedLM Dependent Variable: score
No. Observations: 27247   Method:             REML
No. Groups:       1       Scale:              2.8547
Min. group size:  27247   Log-Likelihood:     -52971.1449
Max. group size:  27247   Converged:          Yes
Mean group size:  27247.0
-----------------------------------------------------
             Coef.  Std.Err.   z    P>|z| [0.025 0.975]
-----------------------------------------------------
Intercept      1.220    1.690  0.722 0.470 -2.091  4.532
distance_TE    -0.000    0.000 -6.844 0.000 -0.000 -0.000
Group Var      2.855
=====================================================
```

**Figure S4:** Scores of TE genes and non-TE genes (left) and correlation of the distance to the closest TE gene with the gene-dispensability score (right) of the *A. thaliana* dataset. TE genes show significantly higher scores than non-TE genes (p ≈ 6E-8; Mann-Whitney U test). Linear mixed modelling shows that there is no relationship between the distance to the closest TE gene and the dispensability score.
