## Additional file 5 for "Reference-based QUantification Of gene Dispensability (QUOD)"

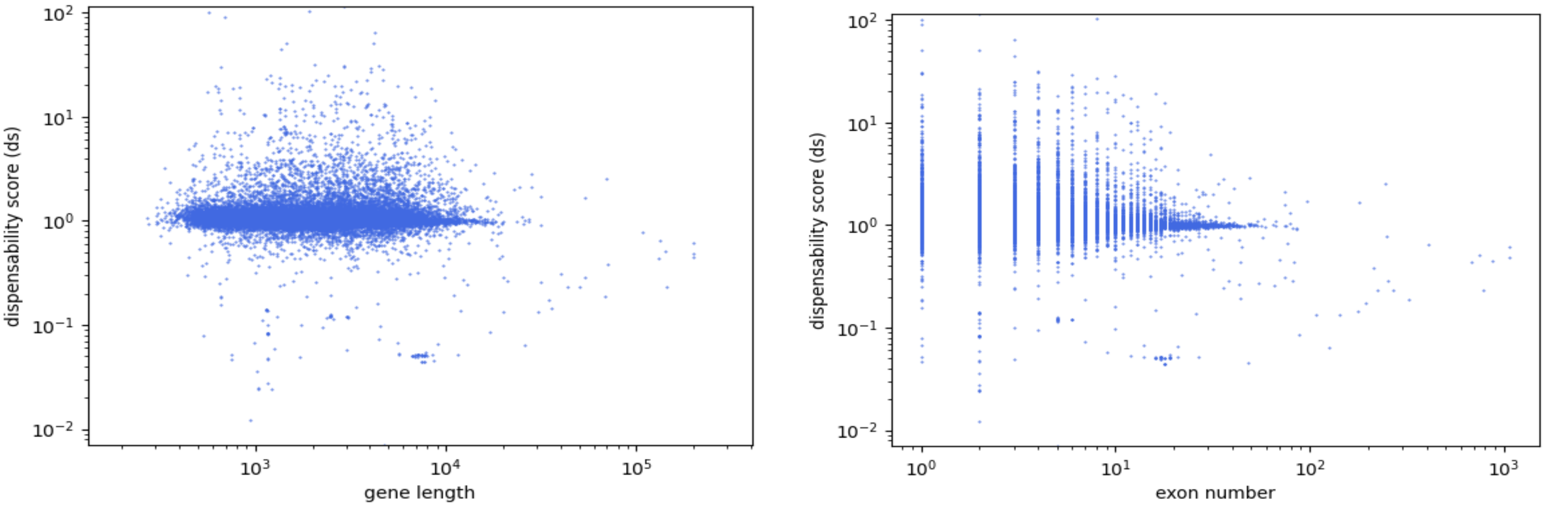


```
        Mixed Linear Model Regression Results
=======================================================
Model:            MixedLM Dependent Variable: score
No. Observations: 30125   Method:             REML
No. Groups:       1       Scale:              2.8722
Min. group size:  30125   Log-Likelihood:     -58657.0610
Max. group size:  30125   Converged:          Yes
Mean group size:  30125.0
-------------------------------------------------------
              Coef.  Std.Err.   z    P>|z| [0.025 0.975]
-------------------------------------------------------
Intercept      1.151    1.695  0.679 0.497 -2.171  4.472
length         0.000    0.000  5.684 0.000  0.000  0.000
exon_number   -0.011    0.002 -6.505 0.000 -0.014 -0.007
Group Var      2.872
=======================================================
```

**Figure S5A:** Correlation of the gene length (left) and exon number (right) with the gene-dispensability score of the *A. thaliana* dataset. Linear mixed modelling shows that there is no relationship between the gene length (Coef. = 0) and the exon number (Coef. = -0.011) and the dispensability score.

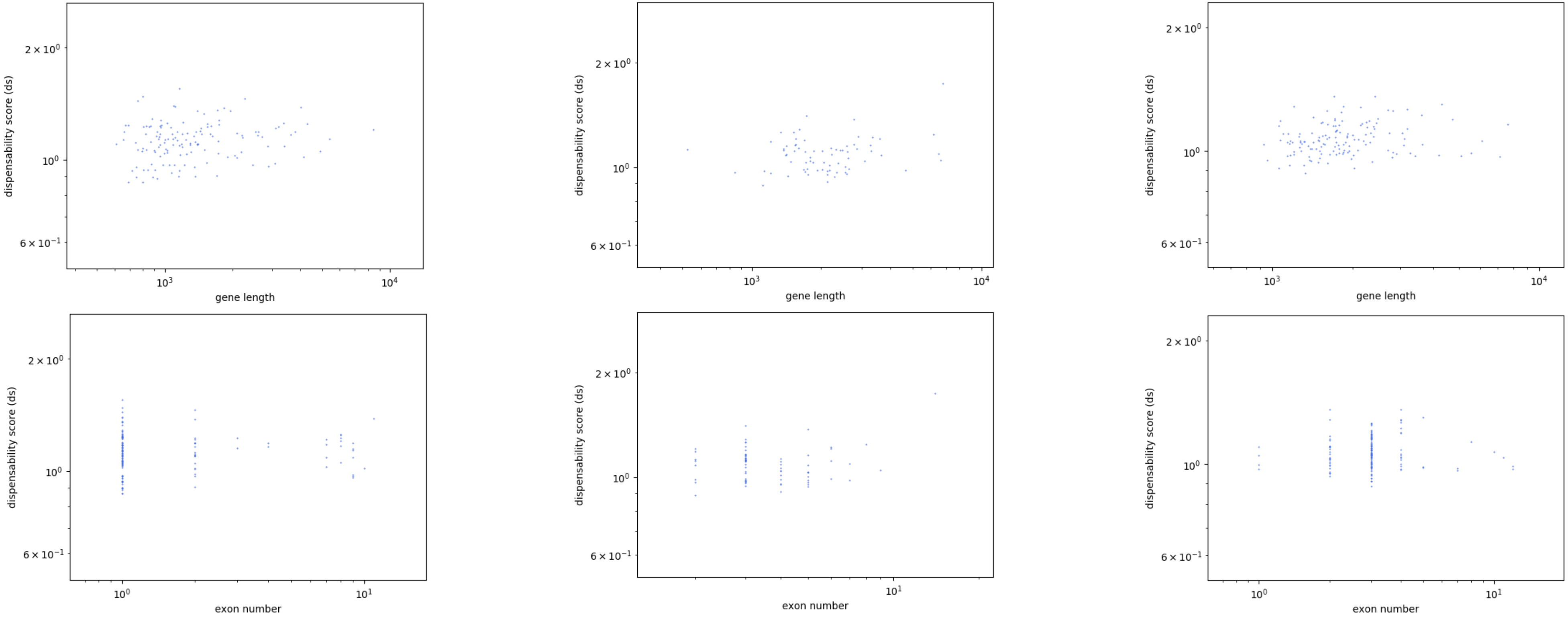


```
Mixed Linear Model Regression Results
================================================
Model:             MixedLM Dependent Variable: ds
No. Observations:  137     Method:             REML
No. Groups:        1       Scale:              0.0181
Min. group size:   137     Log-Likelihood:     64.0370
Max. group size:   137     Converged:          Yes
Mean group size:   137.0
------------------------------------------------------
        Coef.  Std.Err.   z    P>|z| [0.025 0.975]
------------------------------------------------------
Intercept    1.098    0.136  8.077 0.000  0.832  1.365
exon_number -0.010    0.008 -1.368 0.171 -0.025  0.004
length       0.000    0.000  1.913 0.056 -0.000  0.000
Group Var    0.018
================================================
```

```
Mixed Linear Model Regression Results
================================================
Model:             MixedLM Dependent Variable: ds
No. Observations:  72      Method:             REML
No. Groups:        1       Scale:              0.0157
Min. group size:   72      Log-Likelihood:     31.8088
Max. group size:   72      Converged:          Yes
Mean group size:   72.0
------------------------------------------------------
        Coef.  Std.Err.   z    P>|z| [0.025 0.975]
------------------------------------------------------
Intercept   0.996       0.130 7.658 0.000  0.741  1.252
exon_number 0.020       0.015 1.316 0.188 -0.010  0.050
length      0.000       0.000 0.356 0.722 -0.000  0.000
Group Var   0.016 4670288.187
================================================
```

```
Mixed Linear Model Regression Results
================================================
Model:             MixedLM  Dependent Variable:  ds
No. Observations:  140      Method:              REML
No. Groups:        1        Scale:               0.0094
Min. group size:   140      Log-Likelihood:      110.7987
Max. group size:   140      Converged:           Yes
Mean group size:   140.0
------------------------------------------------------
        Coef.    Std.Err.    z    P>|z| [0.025 0.975]
------------------------------------------------------
Intercept    1.067       0.099 10.800 0.000  0.873  1.260
exon_number -0.008       0.006 -1.398 0.162 -0.019  0.003
length       0.000       0.000  1.812 0.070 -0.000  0.000
Group Var    0.009 2454571.831
================================================
```

**Figure S5B:** Correlation of the gene length (top) and exon number (bottom) with the gene-dispensability score of the *A. thaliana* dataset for three different gene families (AP2 (left), WRKY (middle) and MYB (right)). Linear mixed modelling shows that there is no significant effect of the gene length and the exon number on the dispensability score for each gene family, respectively.
