## Additional file 6 for "Reference-based QUantification Of gene Dispensability (QUOD)"

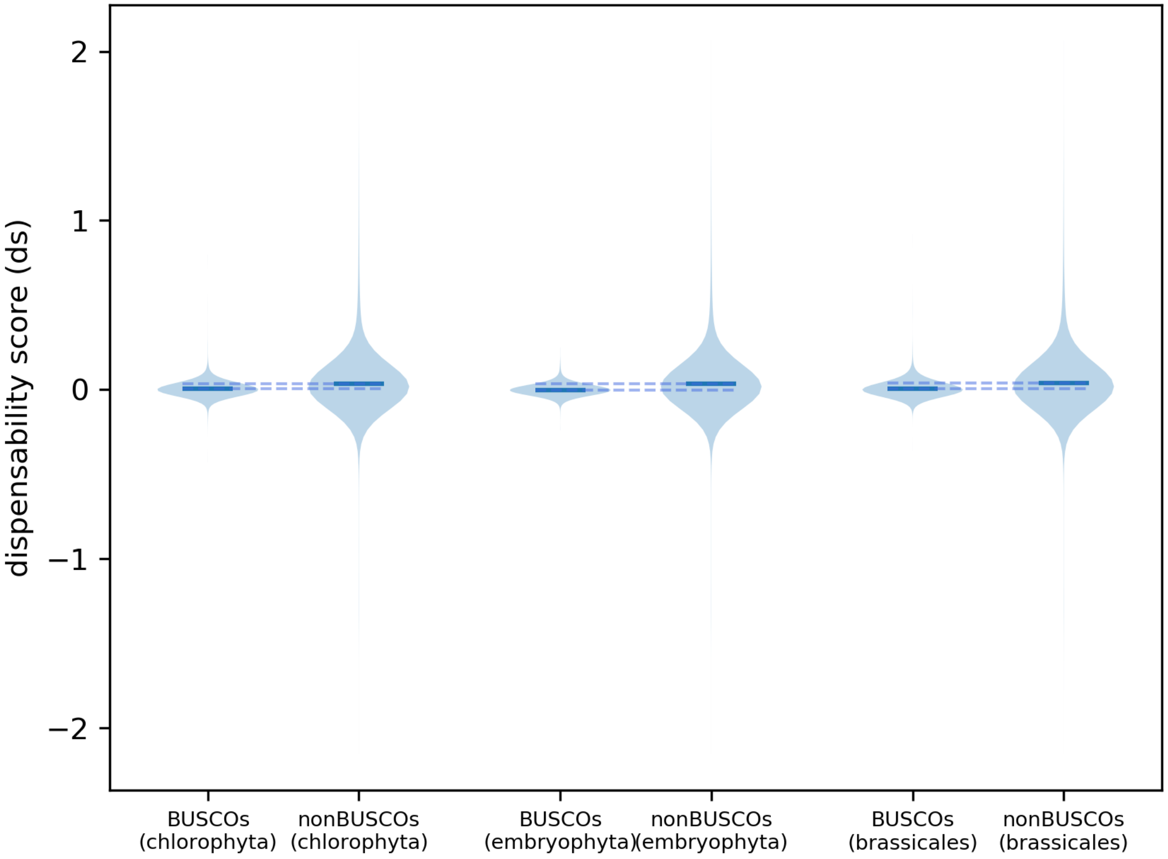


**BUSCO statistics**

| | |
|---|---|
| embryophyta | C:98.4%[S:97.6%,D:0.8%],F:0.2%,M:1.4%, n:1614 |
| brassicales | C:98.3%[S:97.0%,D:1.3%],F:0.1%,M:1.6%, n:4596 |
| chlorophyta | C:91.5%[S:56.9%,D:34.6%],F:0.3%,M:8.2%, n:1519 |

**Mann-Whitney U test**

| | |
|---|---|
| chlorophyta | p ≈ 4E-61 |
| embryophyta | p ≈ 4E-70 |
| brassicales | p ≈ 4E-113 |

**Levene’s test**

| | |
|---|---|
| chlorophyta | $\sigma^2$ (BUSCOs) ≈ 0.021<br>$\sigma^2$ (nonBUSCOs) ≈ 3.590<br>p ≈ 4E-08 |
| embryophyta | $\sigma^2$ (BUSCOs) ≈ 0.005<br>$\sigma^2$ (nonBUSCOs) ≈ 3.497<br>p ≈ 3E-06 |
| brassicales | $\sigma^2$ (BUSCOs) ≈ 0.023<br>$\sigma^2$ (nonBUSCOs) ≈ 3.897<br>p ≈ 1E-14 |

**Figure S6:** Comparison of dispensability scores of BUSCO and non-BUSCO genes using different references (chlorophyta, embryophyta and brassicales). The respective means are represented by the blue lines (dashed lines=extended lines of the respective mean). BUSCO genes show significantly lower scores than non-BUSCO genes for all three reference datasets (Mann-Whitney U test).
Levene’s test was used to test for equal variances. The results show that the variances for all reference datasets differs significantly between BUSCO and non-BUSCO genes. Thus, the deviation of the dispensability score from the respective mean is significantly higher for non-BUSCO genes in comparison to BUSCO genes.
