## Additional file 9 for "Reference-based QUantification Of gene Dispensability (QUOD)"

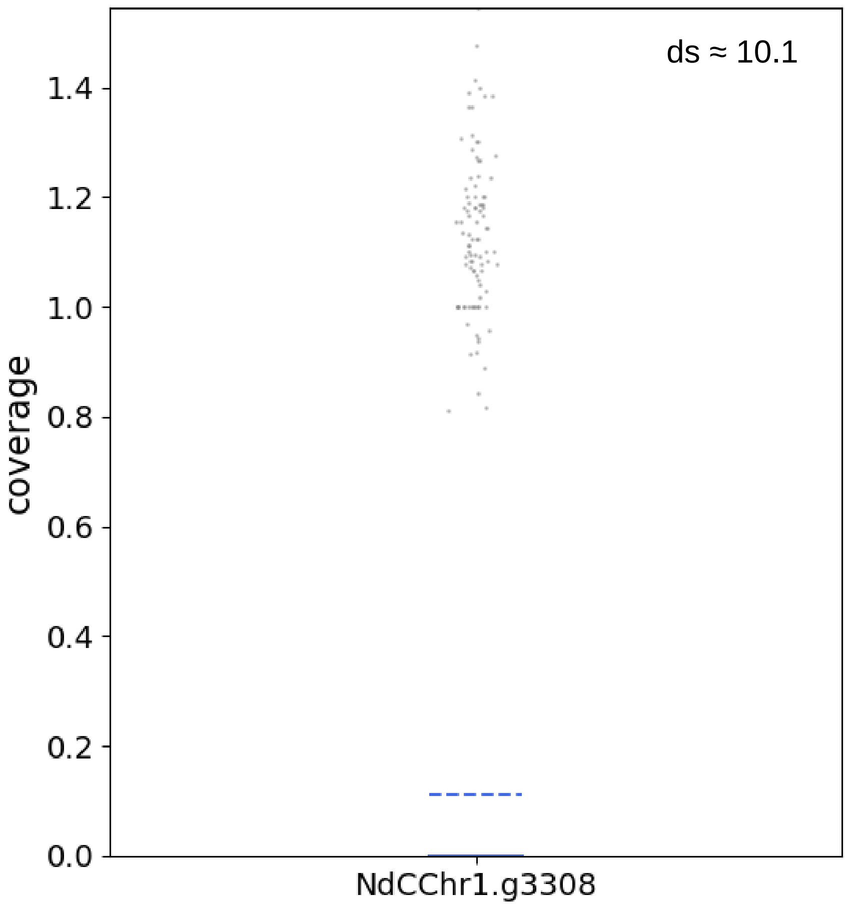


**Figure S9:** Example for lineage specific adaptation. The gene NdCChr1.g3308 (Nd-1 annotation nomenclature) (ds ≈ 10.1) has zero coverage in 870 accessions (≈ 90%) and is annotated as resistance gene mediating resistance against the bacterial pathogen Pseudomonas syringae.
