## Additional file 10 for "Reference-based QUantification Of gene Dispensability (QUOD)"

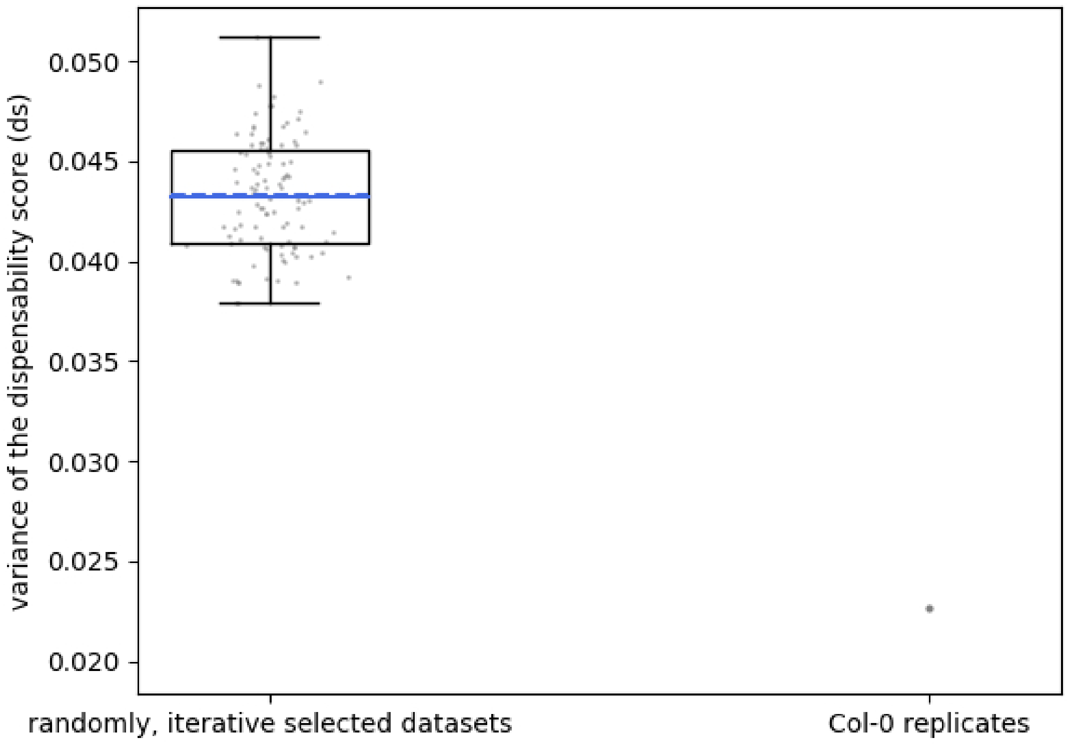


**Figure S10:** A total of 14 genomic datasets of the *A. thaliana* accession Col-0 were used to assess the technical variation between replicates of the same accession. Each dataset was mapped to the TAIR10 reference genome sequence using BWA-MEM. The mappings were then subjected to QUOD expecting a dispensability score close to one for each gene as there should be no variability within the same accession. To test the variation between replicates, we looked at the dispensability score per gene in the dataset containing 14 Col-0 datasets as well in the subsets of all accessions. Therefore, we ran QUOD 100x for 14 randomly selected accessions and once for the Col-0 dataset. We then calculated the variance of all scores for each QUOD run. The Levene's test was applied on the mean scores for each gene of the random subsets compared to the scores for each gene of the Col-0 dataset. The variation of the gene dispensability score distribution of the replicate dataset (one accession) ($\sigma^2 \approx 0.0226$) is significantly lower than the variation between all iteratively, randomly selected subsets of A. thaliana accessions ($\sigma^2 \approx 0.0392$) (Levene's test, $p \approx 4e-19$).
