## Additional file 11 for "Reference-based QUantification Of gene Dispensability (QUOD)"

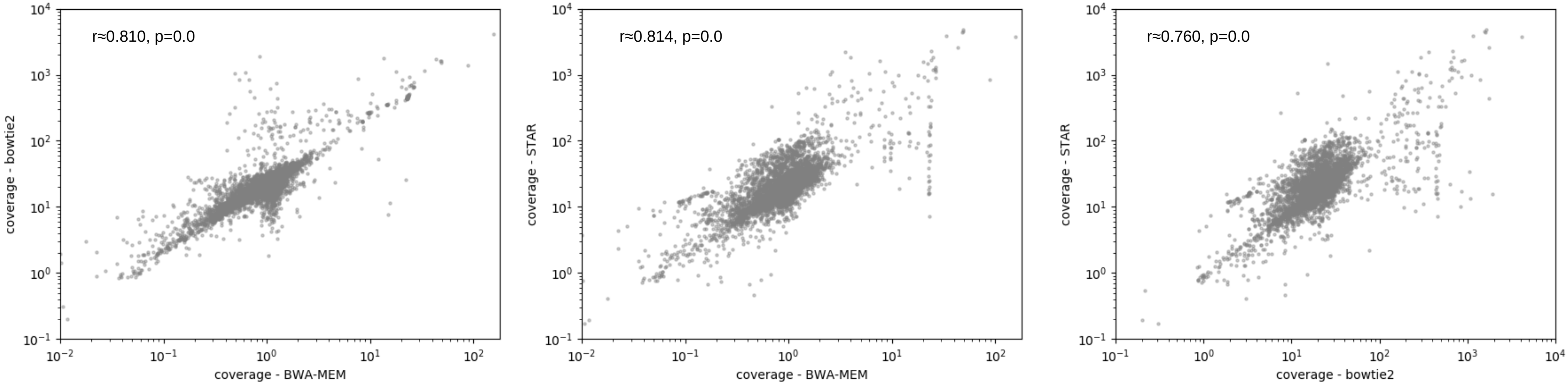


**Figure S11:** Coverage comparison using different read mappers (BWA_MEM, bowtie2, STAR). Spearman correlation coefficient was used to determine the correlation and the significance of the results. The coverages of the genes using different mappers correlate significantly.
